# Z-RNA and the flipside of the SARS Nsp13 helicase

**DOI:** 10.1101/2022.03.03.482810

**Authors:** Alan Herbert, Maria Poptsova

**Affiliations:** InsideOutBio, 42 8th Street, Charlestown, MA; Laboratory of Bioinformatics, Faculty of Computer Science, National Research University Higher School of Economics, 11 Pokrovsky Bulvar, Moscow, Russia 101000

**Keywords:** coronavirus, SARS, MERS, Z-RNA, ZBP1, Necoptosis

## Abstract

We present evidence that the severe acute respiratory syndrome coronavirus (SARS) non-structural protein 13 (Nsp13) modulates the Z-RNA dependent regulated cell death pathways [1]. We show that Z-prone sequences (called flipons [2]) exist in coronavirus and provide a signature (Z-sig) that enables identification of the animal viruses from which the human pathogens arose. We also identify a potential RIP Homology Interaction Motif (RHIM) in the helicase Nsp13 that resembles those present in proteins that initiate Z-RNA-dependent cell death through interactions with the Z-RNA sensor protein ZBP1. These two observations allow us to suggest a model in which Nsp13 down regulates Z-RNA activated innate immunity by two distinct mechanisms. The first involves a novel ATP-independent Z-flipon helicase (flipase) activity in Nsp13 that differs from that of canonical A-RNA helicases. This flipase prevents formation of Z-RNAs that would otherwise activate cell death pathways. The second mechanism likely inhibits the interactions between ZBP1 and the Receptor Interacting Proteins Kinases RIPK1 and RIPK3 by targeting their RHIM domains. Together the described Nsp13 RHIM and flipase activities have the potential to alter the host response to coronaviruses and impact the design of drugs targeting the Nsp13 protein. The Z-sig and RHIM domains may provide a way of identifying previously uncharacterized viruses that are potentially pathogenic for humans.

## Introduction

### Human Coronaviruses

Seven human coronaviruses (hCoV) of the order *Nidovirales* are known to infect humans [3; 4; 5]. These include two alpha-coronaviruses (hCoV-229E (229E)and hCoV-NL63 (NL63)) and five beta-coronaviruses (SARSCoV (SARS1), SARS-CoV-2 (SARS2), MERS-CoV (MERS), hCoV-HKU1 (HKU1), and hCoV-OC43 (OC43)) (Figure 1A). The HKU1, 229E, NL63 and OC43 viruses produce mild symptoms while SARS1, SARS2 and MERS cause severe acute respiratory syndrome (SARS). Much interest has focused on factors that contribute to the morbidity and mortality associated with pathogenic CoV. These include those proteins produced by the virus to counter host responses to the viral double-stranded RNA produced during replication. For example, viral Non-Structural Proteins (NSP) Nsp4a and 4b of MERS suppress induction by double-stranded RNA (dsRNA) of interferon lambda 1 gene expression [6]. A number of other CoV countermeasures interfere with host interferon responses (summarized in [7]). Host factors also increase the severity of disease, including genetic variants that reduce IFN responses and auto-antibodies that inhibit IFN activity [8; 9; 10].

**Figure 1.**
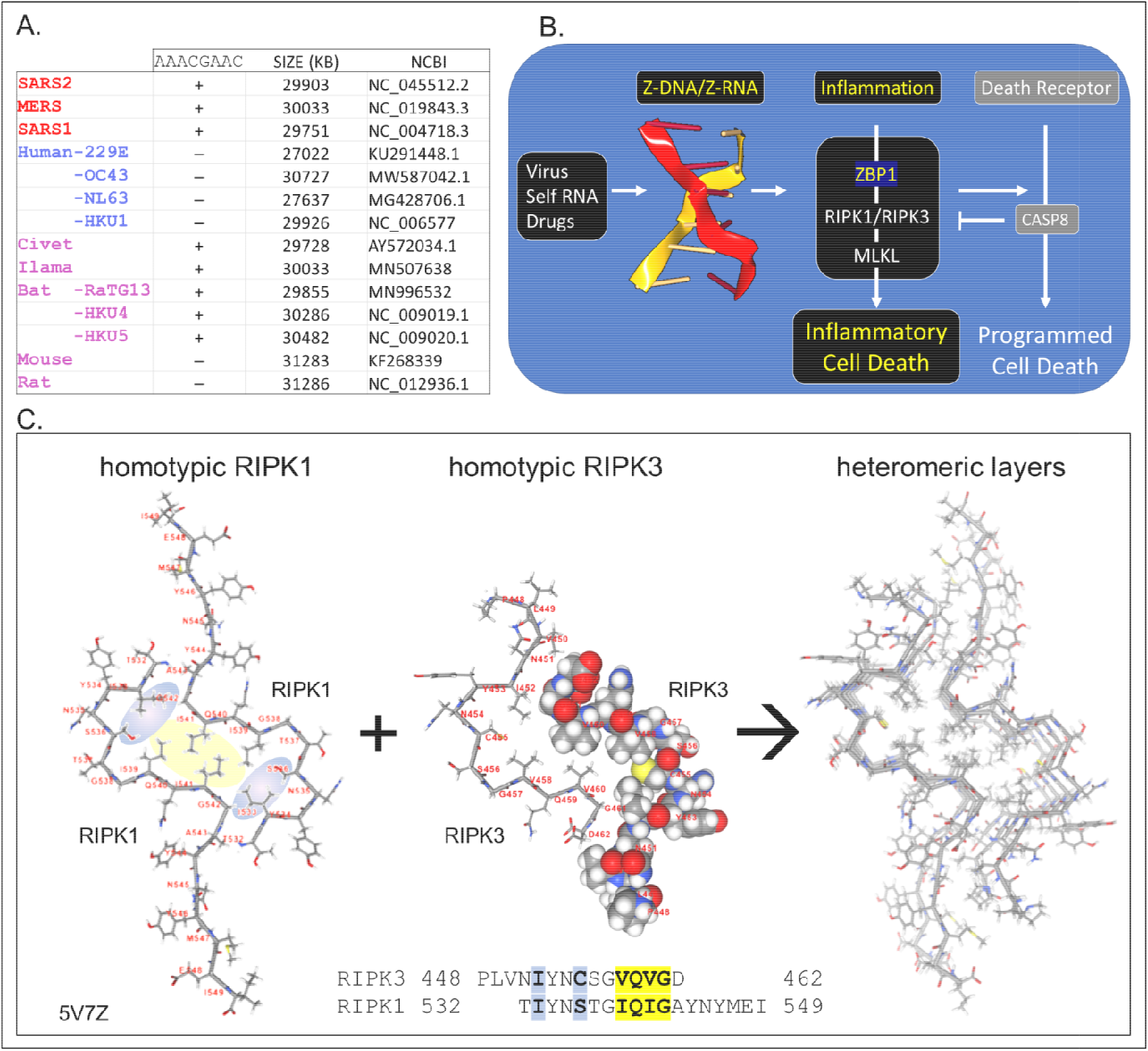
Coronaviruses and human host susceptibility **A**. Coronaviruses (CoV) with the AACGAAC transcription regulatory sequence (TRS) and red labels are associated with severe disease in humans. This TRS is present in CoV from other species but not in all human CoV, especially those that cause mild disease **B**. Host responses to double-stranded Z-RNA viruses can produce inflammatory cell death through the activation of Receptor Interacting Protein Kinases (RIPK) by Z-DNA binding Protein q (ZBP1). The interaction between RIPK and ZBP1 is through their RIP Homotypic Interaction Motifs (RHIM). **C**. Dimers formed from each RHIM protein stack on each other to form a fiber that provides a scaffold for the host response machinery to assemble. The interaction of RIPK1 and RIPK3 is shown with the sequences contributing to the hydrophobic core highlighted in yellow. Blue highlights other residues that stabilize the score. The central figure shows a space-fill representation of one RIPK3 chain present in the dimer.

CoV are long (∼30Kb) positive strand, single-stranded RNA (ssRNA) viruses that replicate themselves within double membrane vesicles through a dsRNA intermediate [4; 11]. The RNA-dependent RNA polymerase (RdRp) involved in genome replication also creates sub-genomic negative strand templates to direct production of structural proteins necessary for virion assembly. The sub-genomic RNAs share the same leader sequence, which is encoded in the 5’ end of the viral genome and joined to a downstream open reading frame (ORF). All the genomic sequence in between that encode the NSPs are excluded. The production of sub-genomic templates depends on transcription regulatory sequences (TRS), such as the AACGAAC sequences present in beta-coronaviruses (Figure 1A). Switching of the RdRp from a downstream TRS to that associated with the leader sequence involves a long-distance RNA interaction [12]. Since replication of the 30Kb CoV genome and the generation of sub-genomic RNAs have the potential to generate sufficient topological stress able to power the flip of right-handed A-RNA to left-handed Z-RNA, we were interested in the role played by the Z-RNA sensor protein ZBP1 (historically called Z-DNA Binding Protein 1) in CoVs infection. After describing Z-DNA and Z-RNA (ZNA) more fully, we propose evidence-based mechanisms through which CoV suppresses Z-RNA dependent inflammatory responses.

### Z-RNA and Flipons

The left-handed Z-DNA and Z-RNA (ZNA) conformations form from their right-handed counterparts of B-DNA and A-RNA by flipping the bases over to produce the characteristic zig-zag backbone of these reverse-twisted, double-stranded nucleic acid conformers [13]. The energy to power the flip can arise from the negative supercoiling produced in the wake of processive enzymes such as helicases and polymerases. In other situations, the topological stress necessary to initiate the ZNA flip is produced by base pairing of entangled nucleic acids, as can occur in stress granules and with defective viral genomes [14]. Sequences that form ZNA under physiological conditions are called flipons. They are often composed of alternating pyrimidine and purine residues, such as (CG)_n_, (CA)_n_ and (UG)_n_. The propensity of flipons to adopt the ZNA conformation can be scored using the ZHUNT3 (ZH3) program [15]. Much of the energy cost in ZNA formation is in the creation of junctions between right-handed and left-handed helices. For Z-RNA, the flip occurs more easily when dsRNA contain basepair mismatches, non-canonical basepairs or unpaired residues at both ends of the Z-forming segment [16].

### Z-RNA and innate immunity

Z-DNA and Z-RNA are recognized in a conformation specific manner by proteins containing the Zα domain, which was first discovered in the dsRNA editing enzyme ADAR1 [17]. The Zα domain is also found in many other proteins, including ZBP1 and virally encoded proteins like vaccinia virus E3 [18] and African swine fever virus [19]. Interaction between Zα proteins regulates cell death pathways [20; 21] (Figure 1B). The pathways are triggered by binding off ZNAs to ZBP1, which then activates RIPK1 and RIPK3 through the RIP Homology Interaction Motifs (RHIMs) shared by all three proteins. RIPK3 once activated by ZBP1 then phosphorylates Mixed Lineage Kinase Domain Like pseudokinase (MLKL) to promote pore formation in cell membranes, producing an inflammatory form of cell death called necroptosis. ZBP1 activation of RIPK1 on the other hand induces Caspase-8-dependent apoptosis, via the adaptor protein FADD and inhibits necroptosis [20; 21].

The RHIM is a ∼40 aa motif, first identified in RIPK1 [22], that contains an (I/V)Q(I/L/V)G sequence at its core [23] (Figure 1C). Structural studies have revealed the nature of how RHIMs interact with each other. The two hydrophobic residues (colored yellow in Figure 1C) of the first RHIM interdigitate with those on a second RHIM to anchor the interaction. The complex is further stabilized by glutamine and other residues (colored in blue) that seal the core and promote the formation of higher order dimensional structures that resemble amyloid fibrils. While RIPK1 and RIPK3 can form homomeric fibrils, they also can stack on top of one another to scaffold assembly of cellular machines that initiate downstream pathways [24; 25; 26].

### Viruses and ZBP1 dependent necroptosis

The ZBP1-initiated cell death pathways plays an important role during infection by the negative RNA stranded influenza virus [27], as well as upon infection with the herpesviruses murine cytomegalovirus (mCMV), human herpes simplex viruses (HHSV) and the vaccinia poxvirus [28]. These viruses have developed a number of strategies to regulate ZBP1-dependent cell death. These include the encoding of ZBP1 homologs, such as the E3 protein, that compete with ZBP1 for ZNAs [28] and proteins with RHIMs that target the ZBP1 and RIPK1/RIPK3 interactions. So far, only RHIMs produced by DNA viruses like the human Herpesviridae (HHV) and mouse cytomegalovirus (CMV) are known to play an important role in virulence [29]. Notably, no RNA virus has yet been shown to encode a RHIM-containing protein.

## Results

### A corona virus-specific Z-flipon signature

We were interested in whether the SARS family coronaviruses might also induce Z-RNA dependent ZBP1 activation and cell death. We used a two-pronged approach. First, we searched for Z-prone sequences in coronavirus using the program ZH3 [30]. We were interested in those sequences that characterize highly pathogenic CoV. Second, we searched for RHIMs in CoV proteins.

### Coronavirus Z-prone sequences

We found that all examined coronaviruses contained sequences that were Z-prone, and that the Z-signature (Z-Sig) for each strain of virus is unique (Figure 1). Using the signature, it is easy to identify the host animal from which the pathogenic human viruses arose. For example, we show that the signatures for Llama coronavirus (CoV) and MERS are identical, as are those for civet CoV and SARS1, as well as those for bat-RaTG13 and SARS2 (Figure 1 Panels A and B). The Z-sigs that characterize each strain are present throughout the genome. Those present in the NSP coding regions of the genome help identify SARS1 and MERS strains, with strong Z-RNA prone regions are also present in the sub-genomic RNAs of both MERS and SARS2.

The ZH3 scoring is based on the energetics of Z-DNA flipping, where a score greater than 600 is sufficient to change flipon conformation under physiological conditions. While not experimentally calibrated for the formation of Z-RNA, the scores observed for the CoVs are far in excess of this value and it is likely hat the sequences in these viruses constitute flipons. With this caveat in mind, we analyzed Z-prone sequences in CoV strains after sequence alignment to the SRAS2 genome, retaining gaps in the output and examine sequences that were present in two or more strains. We also aligned individual CoV gene sequences separately to correct mistakes in the whole genome analysis that arise because of the different sizes in the genome (Figure 3). As expected from the Z-Sigs, the patterns of Z-RNA prone sequences were similar between SARS1 and civet, between SARS2 and Bat-RaTG13 and between MERS and llama-Qatar15.

We analyzed ten of the Z-sig regions identified further. Five Z-sigs were present in genetic regions highly conserved between strains. These encode the Nsp12, Nsp13, Nsp14, S and M proteins. The Nsp13 protein (highlighted in the red box of Figure 3A) is a helicase essential for genome replication and transcription. Its genomic sequences contained the highest Z-Score for SARS1. The score for the equivalent region in SARS2 was lower due to three synonymous mutation in adjacent codons that preserve a hairpin sRNA structure and the encoded peptide PARAR (amino acid (AA) residues 335-339 of SARS2 Nsp13) (Figure 3B). This arginine rich peptide sequence has the potential to bind nucleic acids (figure 2A and B) [31]. The equivalent region of MERS Nsp13 contained a sequence that was less Z-prone and coded for the peptide PA<u>K</u>AR that contains the conservative substitution of lysine (underlined) for arginine. Another highly conserved Z-prone sequence was found in the Nsp12 (RdRp) gene, within the region encoding the NiRAN domain. This domain catalyzes the NMPylation of the strongly conserved N terminus of Nsp9, a modification essential for CoV genomic replication [32]. Although there were nucleotide differences between SARS1 and SARS2, they did not alter the AA sequence. The alternating pyrimidine/purine stem formed by folding the RNA is 11 basepair (bp) in SARS1 rather than 9 bp in SARS2. Another Z-prone sequence was found in Nsp14 and is conserved in strains with different hosts. The Z-prone sequence maps just before the flexible hinge region of the guanine-N7-methyltransferase domain that synthesizes the 5’-terminal cap structure essential for viral mRNA translation [33]. A Z-prone sequence was also present in the region that codes for the S2 ectodomain of the spike protein. The peptide produced includes cysteine 851, which forms a disulfide bond with residue 840 (residues 822 and 833 in SARS), creating a loop that regulates membrane fusion of the virion with the cytoplasmic membrane of the host [34]. Another Z-prone sequence is present in the highly conserved M gene. The segment encodes the membrane protein’s carboxy-terminus AAs that lie just inside the viral envelope. The M-protein interacts with all other structural CoV proteins and is the most abundant protein produced. The protein shapes the virion structure and stabilizes the nucleoprotein (N) protein-RNA complex [35].

**Figure 2.**
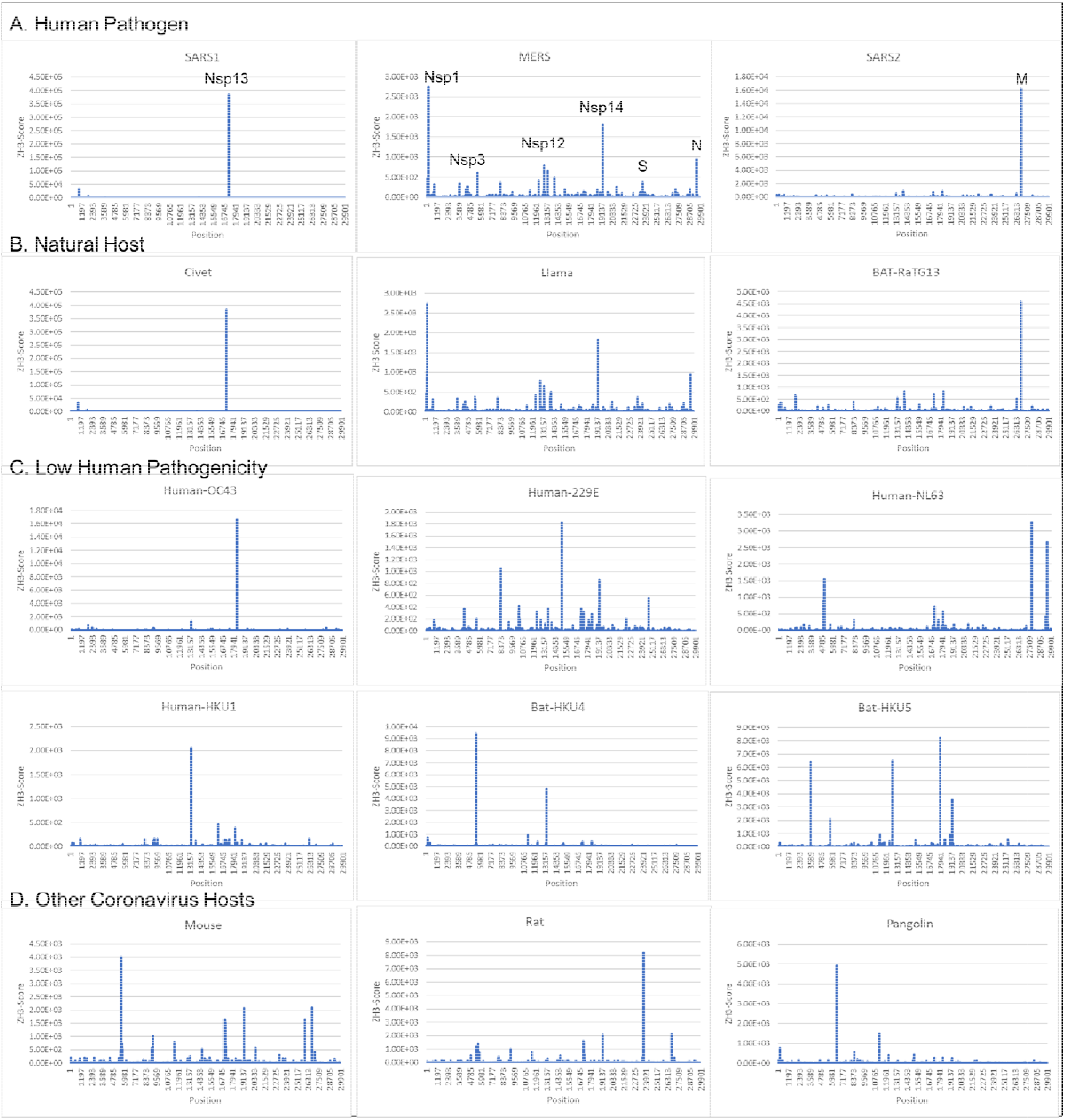
Coronavirus signatures (Z-sigs) based on Z-flipons **A**. Z-sigs for each human pathogenic virus are unique. **B**. Z-sigs match those for the presumed natural hosts for each virus. **C**. The Z-Sigs differ from those for other known coronaviruses **D**. The Z-sigs also vary with the host animal infected. The Z-sig was derived using the ZHUNT3 (ZH3) program [15] that scores the propensity of sequences to flip to the left-handed helical Z-conformation. Sequences with a score greater than 600 are likely to adopt the Z-conformation under physiological conditions. Sequences were aligned using MAFFT prior to analysis [59].

**Figure 3.**
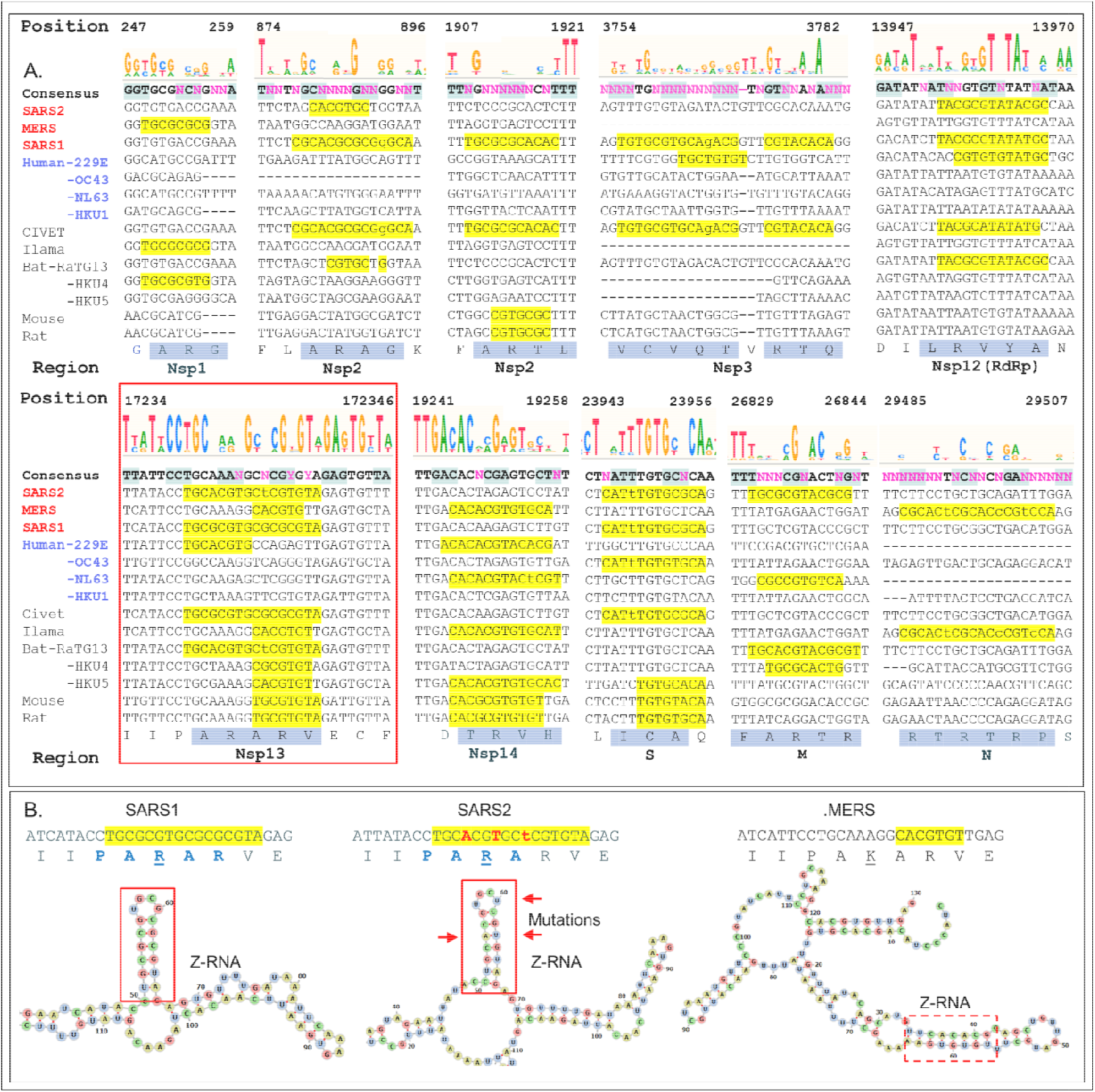
Regions with high ZH3-scores (>600) **A**. The multiple sequence alignment of coronaviruses to SARS2 genome shows potential Z-forming elements in a number of regions of the coronavirus genome. Those encoding Nsp1, Nsp2 and Nsp3 and N proteins contain highly variable regions, while those for the RNA-dependent RNA polymerase (RdRp, Nsp12), the Nsp13 helicase (inside the red box) and the Nsp14 nuclease/methyl-transferase show greater constraint. Lower case letters indicate bases that break the pyrimidine/purine alternation. The amino acid translation of the first and longest nucleotide sequence with a yellow background are highlighted with a blue background. Nucleotide positions are numbered by the SARS2 genome. **B**. Compared to SARS1, the helicase of SAR2 contains three adjacent non-synonymous mutations that conserve the Z-RNA stem and the peptide sequence, but diminish its Z-forming potential. The equivalent MERS sequence has a very low Z-forming potential and encodes a peptide with a lysine (<u>K</u>) in place of an arginine (<u>R</u>). A potential MERS Z-RNA prone element is close-by (within the dashed box).

With regard to other changes in Z-prone sequences, 5 were highly variable between strains. Nsp1 and Nsp2 sequences were quite diverse with some Z-prone sequences encoding an alanine/arginine dimer (AR), suggesting that there regions interact with RNA [31]. Within Nsp1, the Z-prone sequences were present at the amino terminus, a region identified as key to inhibiting the translation of cellular proteins through interactions with the ribosome and critical to a productive viral infection [36]. Nsp1 is thus subject to selective pressures that varies by host. Nsp2 contains two Z-prone sites. The protein is known for its poor sequence conservation with only the zinc finger domain preserved [37]. Another Z-sig is present in Nsp3, which is quite a long gene that also varies in composition between CoV strains [38] The Z-prone sequence in SARS1 Nsp3 encodes part of the macro-1 domain that is involved in de-mono-and de-poly-ADP-ribosylation. The SARS2 sequence is VCV<u>D</u>TVRT<u>N</u> while the SARS1 sequence VCV<u>Q</u>TVRT<u>Q</u>, differing by the two residues underlined. These amino acid changes do not directly involve the active site. At the nucleotide level, SARS1 RNA folds on itself to form a 7 bp Z-RNA stem whereas SARS2 RNA does not. Z-prone sequences within the N-protein gene are present in MERS strains. This gene codes for the nucleoprotein involved in packaging the viral RNA into the virion. The Z-sig encode part of the unstructured C-terminus of the protein, beyond the domain critical for RNA packaging that has been characterized in crystallographic studies [39]. The Z-sig encoded residues RTRTR (395-399) do have an alternating motif similar to the KAKAK motif that is specific for Z-RNA [40], potentially allowing interacting with Z-prone sequences in that genome, possibly phasing the sites in the genome to which the N-protein binds.

### A putative RHIM associated with coronaviruses severely pathogenic in humans

In assessing the potential for Z-RNA dependent responses, we searched for viral proteins containing a RHIM that could modulate cell death pathways through interactions such as those illustrated in Figure 1. A multiple sequence alignment showed that Nsp13 in MERS, SARS1 and SARS2 have similar features to known functional RHIMs that are absent in the 229E, OC43, NL63 and HKU1 CoVs (Figure 4A). The residues are colored as in Figure 1C with the core motif in yellow and the other residues that seal the fold in blue. The gray box highlights the labels for previously identified RHIMs. The potential RHIM identified in the nonsense mediated decay associated (NMD) kinase SMG1 [41] is currently undocumented. Other potential RHIMs in Nsp14 (<u>V</u>GT<u>N</u>LP<u>LQLGF</u>), in SMG1 (<u>K</u>AV<u>E</u>HN<u>IQIG</u>K) and UPF1 (<u>R</u>PM<u>F</u>FY<u>VQLG</u>Q) did not align well with known RHIM sequences, but warrant further investigation to test their functionality.

**Figure 4.**
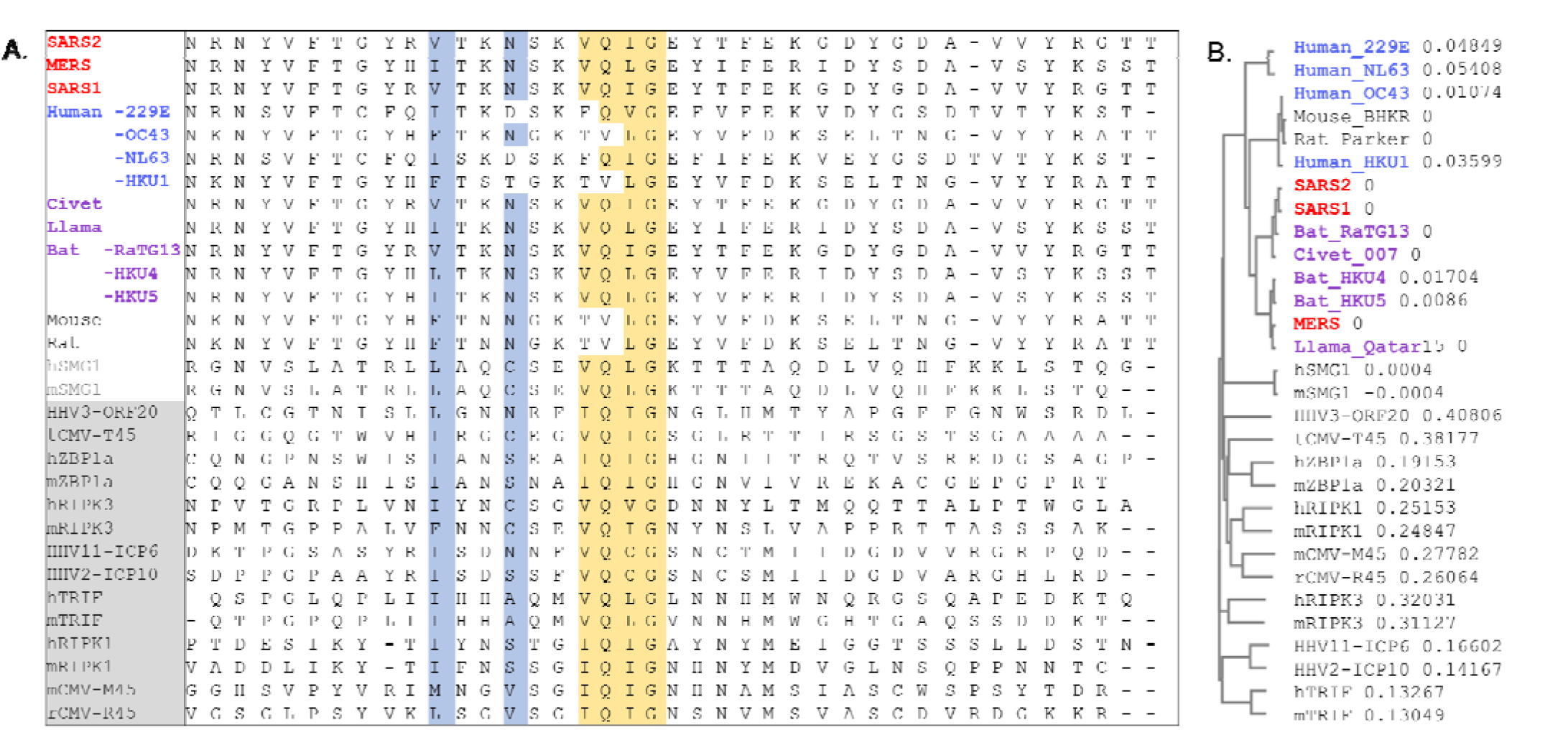
RIP homotypic Interaction motif. **A**. CoV Nsp13 protein sequences from different strains that contain the RHIM (I/V)Q(I/L/V)G residues (in yellow) were aligned with known RHIMs from other viral and host proteins (labels in the gray box). SMG1 was identified here but is not yet a validated RHIM. Residues are colored as in Figure 1. **B**. Cladogram based on distance to show the relationship of proteins. (HHV: human herpes virus; CMV: cytomegalovirus; ZBP1: Z-DNA binding Protein 1; RIPK; Receptor Interacting Protein Kinase; TRIF: Toll like Receptor Adaptor Molecule; SMG: SMG1 nonsense mediated mRNA decay associated PI3K related kinase; h: human; m: mouse; r: rat; t: tupaiid)

### The structure of Nsp13 helicase

Both strategies of looking for Z-prone sequences and RHIMs focused our attention on the Nsp13 helicase and its potential novel role as a Z-flipon helicase (flipase). Nsp13 unwinds both DNA and RNA in the 5’->3’ direction. It is a member of the helicase superfamily 1B (SF1B) [42]. Along with Nsp12 and other CoV NSP proteins, Nsp13 forms the viral replication-transcription complex. High resolution crystallographic and electron-microscopy structures of Nsp13 have recently been published and show that a second Nsp13 cooperates with the first to enhance translocation of Nsp13 along the CoV genomic RNA [43; 44; 45]. The Nsp13 helicase has two canonical RecA domains between which ATP is bound and through which single-stranded RNA egresses from the complex. Nsp13 also has three N-terminal domains that are unique to nidovirus helicases, with a stalk domain connecting a zinc-binding (ZincBD) domain to the 1B domain (Figure 5).

**Figure 5.**
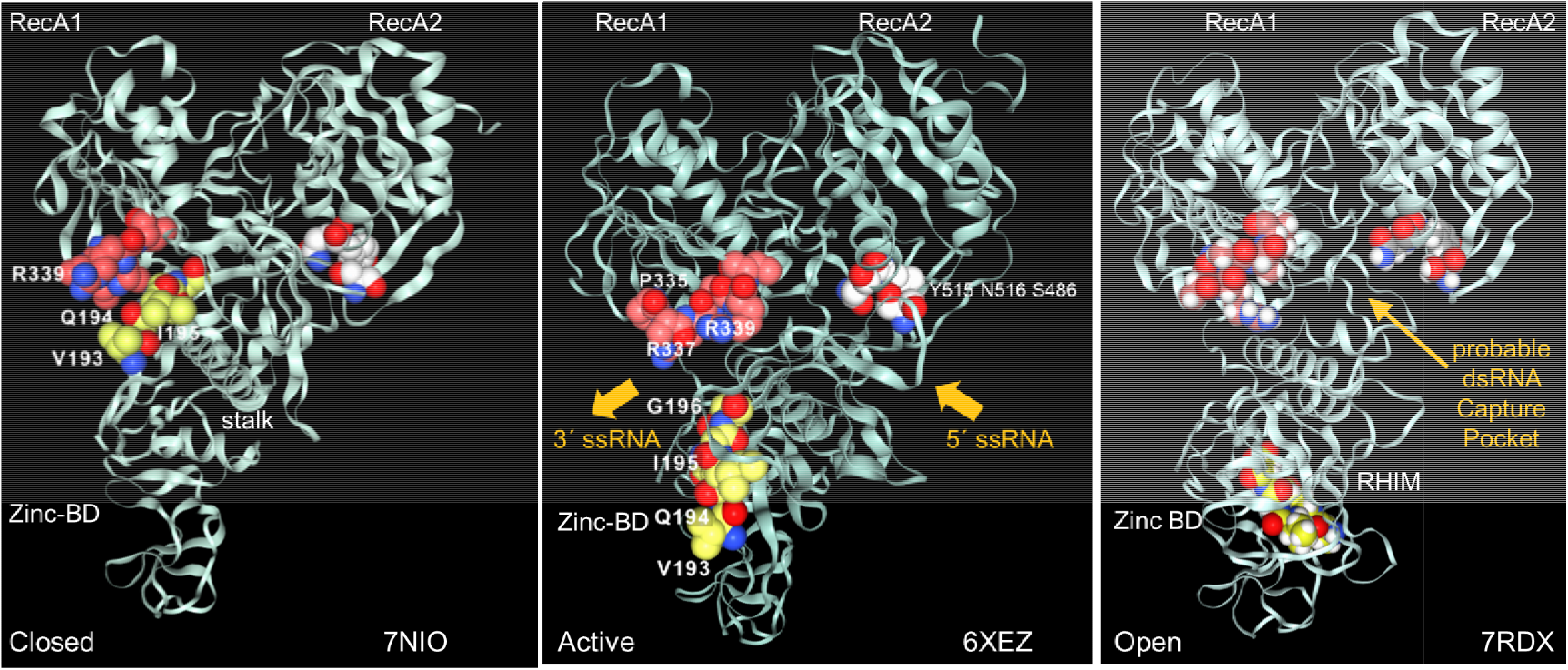
The interaction between the Nsp13 RHIM (residues 193-6) and PARAR (residues 335-9) are conformation dependent. **A**. In the ATP-free apo form of Nsp13, the RHIM (yellow space fill carbons) contacts PARAR (crimson space fill carbons) **B**. In the active state, RHIM and PARAR separate, opening the 5’ end of the single-strand RNA channel, that has the 3’end marked by N516 (white space fill carbons). **C**. In the open complex, the RHIM separates further from the RecA2 domains to create a cavity that is potentially large enough to accommodate dsRNA. This opening is associated with a rotation of the Zinc binding domain (ZincBD) relative to N516. PARAR also rotates, changing the position of R337 and R339. The structures are from PDB files 7NIO [43], 6XEZ [44] and 7RDX [45] as labeled, with images rendered using the NGLViewer [56].

The Nsp13 RHIM is present in the 1B domain, which bridges the RecA1 and RecA2 domains. Rather surprisingly, the peptide PARA encoded by the Z-prone sequence that is in the RecA1 domain interacts with the RHIM when the Nsp13 is in the ATP-free closed conformation (Figure 5A). In the active conformation, the RHIM and PARA segments separate to open a channel through which the ssRNA passes (Figure 5B). Further separation of the 1B domain from the RecA2 domain renders Nsp13 unable to translocate on RNA [45]. The cavity created by this separation appears flexible enough to accommodate a dsRNA structure (Figure 5C).

The Nsp13 cavity shown in Figure 5C is lined by the residues known to bind ssRNA (shown by white space fill carbons in figure 6A and 6B), both to the phosphate backbone and also in a base-specific manner. Into the newly formed cavity projects the PARA peptide, with R337 and R339 forming a hook (Figure 6A). Also, tyrosine Y205, along with W178, faces the interior of the cavity (Figure 6B). Potentially Y205 and W178 could engage with each other in an edge to face configuration similar to that of Y177 and W195 in the Zα domain to form a platen for the conformation-specific recognition of Z-RNA. These findings support a new role for Nsp13 as a flipase that is described further in the discussion below.

**Figure 6.**
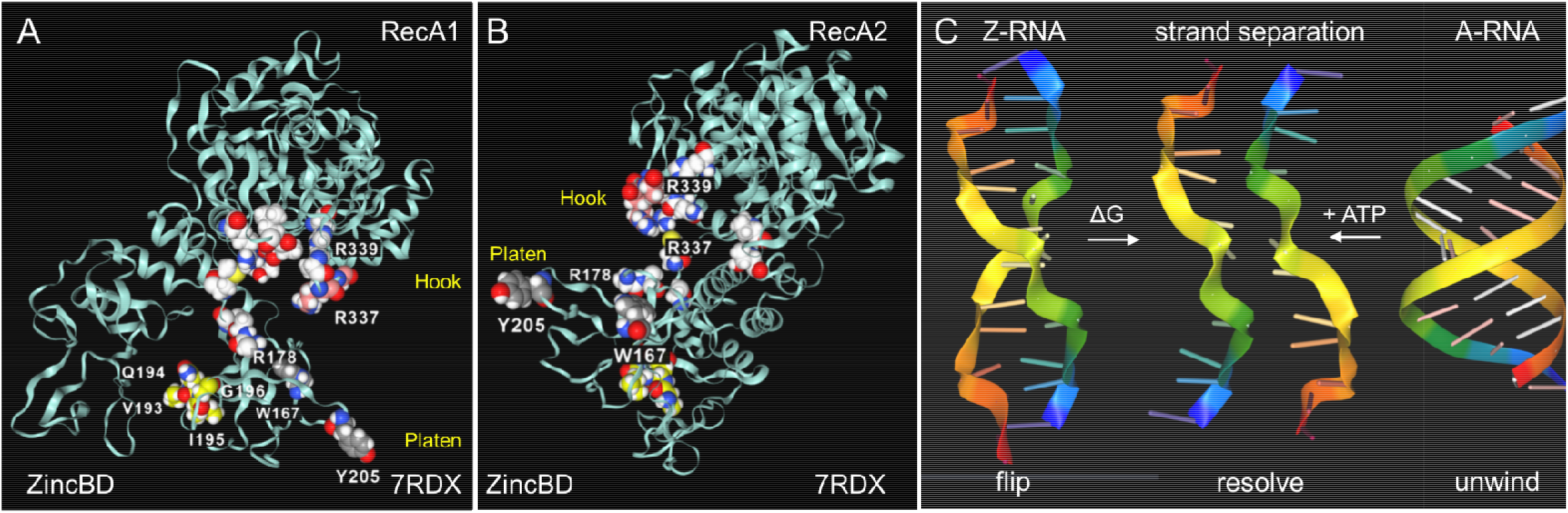
Nsp13 opens to expose a hook and platen for binding dsRNA. **A**. The arginine rich PARAR hook (crimson carbons) and the tyrosine (Y205, grey carbons) platen create a surface for docking dsRNA. Tryptophan (W168)(blue carbons) has the potential to orientate Y205 through and edge to face interaction and create a Z-RNA specific recognition element like that present in the Zα domain and confers specificity for binding to the left-handed helix [61]. In this open orientation, the RHIM (yellow carbons) is free to engage other RHIM proteins. The space fill with white carbons show the residues identified as making base-specific contacts with single-stranded RNA (ssRNA) in the active conformation of Figure 3B (residues 178, 179, 230, 233 in domain 1B, residues 311, 335,361, 362:E,363, 390, 408, 410:E in RecA1). **B**. The view of the Nsp13 molecule from the opposite side to that shown in the first panel. **C**. In addition to the ATP-dependent helicase activity, Nsp13 has the potential to capture ssRNA when Z-RNA flips to and from A-RNA. The higher energy Z-RNA powers the strand-separation, providing the ΔG needed to fuel the Z-flipon helicase activity we refer to as a Z-flipase.

## Discussion

### CoV Z-RNA prone sequences

Our analysis reveals a number of Z-RNA prone sequences (Z-sigs) in human CoV that can be used as fingerprints to identify closely related stains to trace their zoonotic origins. The approach is facile without the need for deep sequence analysis or alignment to confirm the relationship and is quite robust to errors in such procedures.

The plots in figure 2 show that potential flipons can occur in many locations within the viral genome. The presence often is associated with alanine/arginine dimers that are encoded by the nucleotide sequence GC(G/A)CG(C/T) and likely reflect in part the essential role of these residues in highly conserved interactions of the virus with RNA [31]. Besides serving as codons, Z-sigs could potentially act as flipons. For example, the flip to Z-RNA could relieve the topological stress generated when both the positive and negative strands of the genome are transcribed by viral polymerases, or during template switching, when the genome loops back to align the leader sequence with more distant regions to produce sub-genomic negative strand RNAs. Alternatively, Z-RNA formation in Z-prone sequences may allow the virus to trap proteins involved in host defense and inhibit their activation of cell-death pathways. Likewise, defective viral genomes, where the Z-RNA flip is powered by base-pairing of entangled RNAs, may act as decoys to derail host responses. Z-prone sequences also can occur in more variable regions of the genome, enabling the evolutionary adaptation of the virus to its host, either through amino acid changes that improve protein function or through the flipon mediated retargeting of the host ribonucleoprotein machinery involved in translation in much the same way as the Z-Box of Alu repeat elements regulate ribosomal assembly [13].

### The CoV host range and RHIMs

Of note is that some hosts lack functional Z-RNA dependent inflammatory cell death pathways, so there is less constraint on the available sequence space they can exploit [46]. However, the Z-prone sequences that evolve in these viruses may impair their ability to infect a new species that have intact ZBP1-dependent cell death pathways. Human represents one such challenging host for zoonotic viruses to colonize. The acquisition of a RHIM domain to prevent Z-RNA dependent outcomes may represent a key adaptive strategy that allows SARS1, SARS2 and MERS strains to productively infect humans. The alternative of reducing the number of viral Z-prone sequences requires retooling of the existing protein machinery and many more changes. In the example of SARS2, the Z-forming potential of the SARS1 Nsp3 Z-sig is reduced by non-synonymous mutations, leaving the encoded peptide the same (Figure 3B). It is unknown whether this decrease in the potential to form Z-RNA increased the potential for SARS2 to become pandemic, but viruses like Epstein-Barr virus show selection against Z-DNA forming sequences relative to other HHVs [47].

### Novel Nsp13 flipon helicase activity

Our structural analysis of the potential Nsp13 RHIM raised the possibility that protein has a non-canonical helicase activity that exploits the strand separation occurring during the transition of A-RNA to Z-RNA (Figure 6C). In this case, the unwinding of dsRNA by the flipase can be powered by the free energy stored in Z-RNA rather than depending like a classical A-RNA helicase does on the energy released by ATP hydrolysis (Figure 6C). The activity of the flipase can be further enhanced by base-specific binding to the alternating pyrimidine/purine ssRNA motifs present in Z-prone sequences, like those present in the CAUGU substrate used by Chen et al in their Nsp13 structural studies. [45]. By sequestering such sequences, Nsp13 would impede the formation of the Z-RNA, inhibiting activation of ZBP1 initiated pathways.

It also appears from the structural analysis shown in Figure 5 that engagement of Nsp13 by Z-prone sequences exposes the proposed Nsp13 RHIM. The motif is on the opposite face of the 1B domain to the one that binds RNA (Figure 3C). Once sprung loose, the RHIM is free to contact and modulate the activity of ZBP1, RIPK1 and RIPK3 (Figure 1B), much as the RHIMs from HHV-ICP6, mCMV-M45 and varicella virus HHV3-ORF20 RHIMs do. All of these known viral RHIMS promote formation of heteromeric amyloids [48]. The higher order structures of each viral fibril differs, likely reflecting the impact of residues outside the central core (Figure 1C and 4A). While M45 preferentially interacts with RIPK3 and ZBP1 over RIPK1, ORF20 primarily sequesters ZBP1 [49; 50]. These differences alter the functional outcomes. M45 inhibits tumor necrosis factor induced necroptosis and other cell death pathways, while ORF20 diminishes only ZBP1 dependent apoptosis.

It is also possible that the RHIM domain of Nsp13 targets other cellular proteins. The RHIM bearing TRIF protein (toll like receptor adaptor molecule 1 encoded by TICAM1) is directly bound by the dsRNA sensor TLR3 but is indirectly recruited to lipopolysaccharide binding TLR4 via the bridging adaptor TRAM [51]. Mice lacking TRIF are highly susceptible to SARS1 infection and show reduced lung function, increased lung pathology, higher viral titers and increased mortality. These outcomes support the possibility that Nsp13 interacts directly with TRIF and recapitulate many features of human disease [52]. Another potential RHIM interaction is with SMG1, a host protein that regulates the onset of NMD by phosphorylating for Upf1 protein. CoV are highly susceptible to NMD due to their dependence on and the nature of the sub-genomic RNAs [53]. The vulnerability arises because only the first ORF of sub-genomic transcripts is translated. The rest of the message is seen by the host as a long 3’UTR of the kind susceptible to NMD [54]. Engaging the potential SMG1 RHIM could be one way the virus inhibits NMD and ensures virion components are synthesized in the quantities required.

In summary, we propose that Nsp13 is able to prevent ZBP1 activation of cell death pathways in two different ways. First, the helicase prevents the Z-RNA formation required to engage ZBP1 and second, the RHIM domain inhibits activation of cell death pathways by ZBP1 (Figures 1B and 4A). Potentially the Nsp13 RHIM also regulate activation of inflammatory responses by TRIF and prevent initiation of NMD by SMG1.

### Disease Implications

At this stage, we only know of the possibilities and hope that the above hypotheses will provide a framework for further investigation of Nsp13 effects on Z-RNA dependent cell death in properly qualified BSL3 laboratories. If it is found that pathogenic CoVs do indeed produce Z-RNA (as the ZH3 algorithm predicts) and that the RHIM in Nsp13 regulates ZBP1, RIPK3, RIPK1, TRIF or SMG1 signaling, then we suggest that effects will depend on the stage of viral infection. At early stages, suppressing Z-RNA formation and cell death/NMD would enable increased viral replication while diminishing severity of the initial infection. Here the recognition of Z-RNA prone sequences by Nsp13 flipase then blocks the viral polymerase until the two RNA strands are fully separated by the classical Nsp13 helicase activity. At later stages of infection, the dsRNA tangles formed as defective viral genomes accumulate may create Nsp13 clusters, leading to the formation of amyloid filaments, then inflammatory cell death as recently reported for *in vitro* and *in vivo* models of SARS2 infection[55].

### Coronavirus and Human Pathology

From our analysis, it seems that the RHIM domain is a requirement for CoV for the successful jump the virus made from its natural host to humans, although not a sufficient one as other mutations that enable engagement of human cell surface receptors are required. The combination in Nsp13 of the RHIM domain with a Z-flipase so far appears unique among human viruses and may account for the extreme virulence of a subset of CoV. The presence of RHIMs in other viruses that are currently poorly characterized may identify them as potential pathogens before they become pandemic in humans, allowing implementation of pre-emptive measures to counter human infections.

The susceptibility of humans to SARS and other novel viruses may also depend on each individual’s genome as there is a high frequency of gene variants that impact the necroptotic machinery [46]. The loss of function alleles reported may leave some individuals more vulnerable to CoV infection and enable persistent viral infections that lead to chronic disease outcomes. Genetic data from cases of long covid would allow identification of any suck risk variants.

While Nsp13 inhibitors that target the ATP-binding site are likely to be effective early in CoV infection by inhibiting the classical helicase function of the enzyme, only those compounds that lock the enzyme in a conformational state unable to engage Z-RNA are likely to be effective against the Z-flipase activity.

## Methods

The Z-forming potential of the Coronavirus sequences were assessed using ZHUNT3 using the following command line “zhunt3nt 12 6 24” [30]. The structures PDB files 7NIO [43], 6XEZ [44] and 7RDX [45] were visualized using the open access NGLViewer [56]. Amino acid alignments and the Phylogram were prepared using MUSCLE [57] using the following NCBI sequences for MERS (NC_019843.3), SARS1 (NC_004718.3), SARS2 (NC_045512.2), Hum_CoV-229E (KU291448.1), CoV-OC43 (MW587042.1), CoV_NL63 (MG428706.1), CoV-HKU1 (NC_006577), Bat RaTG13 (MN996532), Civet_007 (AY572034.1) and Llama (MN507638), BAT_HKU4 (NC_009019.1), BAT_HKU5 (NC_009020.1), Murine_CoV_MHV/BHKR (KF268339), RAT_Parker (NC_012936.1) and Pangolin (OM009282.1); Tupaiid herpesvirus 1 tCMV (NC_002794.1). The RHIM in the SMG1 was identified using the MEME Suite [58], using the 10 residues incorporating the colored residues shown in Figure 4 as a seed. Sequence alignments were performed using the MUSCLE [57] CLUSTAL OMEGA algorithm for protein sequences and the CLADOGRAM shown in Figure 4B (accessed on March 24^th^, 2022 at https://www.ebi.ac.uk/Tools/msa/clustalo/ and MAFFT V7 for Coronavirus sequence alignment, using the option to preserve gaps with the others left as the default and accessed online on March 24^th^, 2022at https://mafft.cbrc.jp/alignment/server/ [59] and motifs produced using the free version of SNAPGene. Structure images were rendered using the NGLViewer [56] and RNAfold was used to render the RNA images shown in Figure 3B [60].

## Author Contributions

AH is an international supervisor for the HSE Bioinformatics Laboratory run by MP. AH conceptualized and wrote the paper with suggestions and edits from MP and AH prepared the figures.

## Acknowledgements

We thank Sid Balachandran for suggesting the possibility of RHIM domains in SARS2 and for helpful edits to the manuscript. We thank Alexander Shein who works with M.P. and identified the presence of a strong Z-forming sequence in SARS1

## Conflict of Interest

Author AH is the founder of InsideOutBio, a company that works in the field of immuno-oncology. All the authors declare that the research was conducted in the absence of any commercial or financial relationships that could be construed as a potential conflict of interest.

## Notes

### Competing Interest Statement

The authors have declared no competing interest.

### Summary of Updates

The analysis is more extensive and the conclusions are better evidenced.

